# Shared evolutionary path in social microbiomes

**DOI:** 10.1101/2023.02.07.527449

**Authors:** Nelson Frazão, Isabel Gordo

## Abstract

Social networks can influence the ecology of gut bacteria, shaping the species composition of the gut microbiome in humans and other animals. Gut commensals evolve and can adapt at a rapid pace when colonizing healthy hosts. Here, we aimed at assessing the impact of host-to-host bacterial transmission on *Escherichia coli* evolution in the mammalian gut. Using an *in vivo* experimental evolution approach in mice, we found a transmission rate of 7% (±3% 2SE) of *E. coli* cells per day between hosts inhabiting the same household. Consistent with the predictions of a simple population genetics model of mutation-selection-migration, the level of shared events resulting from within host evolution is greatly enhanced in co-housed mice, showing that hosts undergoing the same diet and habit are not only expected to have similar microbiome species compositions but also similar microbiome evolutionary dynamics. Furthermore, we estimated the rate of mutation accumulation of *E. coli* to be 2.9×10^−3^ (±5×10^−4^ 2SE) mutations/genome/generation, irrespective of the social context of the regime. Our results reveal the impact of bacterial migration across hosts in shaping the adaptive evolution of new strains colonizing gut microbiomes.

## Introduction

Individuals that share the same space are known to harbor more similar microbiota compositions than individuals that do not. This indicates that microbial migration between hosts, which is increased when living in the same household, is an important factor structuring species diversity in the microbiome (Johnson and Clabots 2006; Johnson et al. 2008; Siranosian et al. 2021). Indeed, it has been shown that co-housing of mice, which are co-prophagic, can reduce the diversity of the microbiota species composition among hosts. Co-housing is also a common practice aimed at reducing microbiota variation in many studies of host immune phenotypes (Ericsson and Franklin 2015).

Emerging data in both mice and humans shows that significant evolutionary change can occur within strains of each microbiota species (Garud and Pollard 2020). In germ-free mice colonized with either a single or multiple strains of commensal bacteria (Li et al. 2015; Barroso-Batista, Pedro, Sales-Dias, Catarina J. G. Pinto, et al. 2020; Yilmaz et al. 2021) or in mice with a native microbiota (Barroso-Batista et al. 2014; Lescat et al. 2017; Frazão et al. 2019; Frazão et al. 2022), evolutionary change has been observed within days, weeks or months. Time series data of human metagenomes also show that several adaptive evolutionary events can occur within months (Garud et al. 2019; Zhao et al. 2019).

To begin unraveling how bacterial transmission across hosts may affect patterns of molecular evolution of their gut microbes, population genetics theory of metapopulations, where a deme mimics a host, can be useful (Pannell and Charlesworth 1999; Booker et al. 2021). Theoretical models of adaptation in the context of metapopulations predict that the rate of adaptive evolution of a clonal population (*i*.*e*., of bacteria) resulting from the accumulation of new beneficial mutations should be affected by the amount of migration/transmission between demes (*i*.*e*., individual hosts). In particular, for a given level of migration, the rate of adaptation should be increased relative to when no migration occurs (Gordo and Campos 2006; Yeaman and Whitlock 2011). The models typically consider a single microbial species and therefore ignore the multispecies complexity characteristic of the gut ecosystem. However, their predictions should be robust for ecosystem level complexities under conditions where the strength of intra-specific competition is much higher than that of inter-species competition. Such conditions are observed when ecological models aiming at explaining the diversity and stability of microbiomes, like the generalized Lotka-Voltera, are fitted to 16s RNA data (Coyte et al. 2015).

Here we study how the evolutionary path of a new lineage colonizing the mouse gut is influenced by its host social environment. We use *in vivo* experimental evolution to test the following hypotheses: i) that co-housing increases the rate of evolution of commensal bacteria, and ii) that high migration rates lead to an increase in the number of shared evolutionary events accumulating across hosts. We test this hypothesis for the two key processes of evolution: horizontal gene transfer (HGT) and mutation accumulation. Under HGT, we expect that adaptive events occurring in only one mouse will transmit rapidly to the entire mouse metapopulation. Under mutation accumulation, we expect higher allele sharing in a social regime if adaptive evolution within the host is marked by intense competition between clones carrying different adaptive mutations. In fact, when clonal interference is pervasive within hosts, which live in the same environment (*e*.*g*., same diet), migration/transmission may help the bacterial clones which carry the highest combination of beneficial alleles to spread more rapidly across hosts.

## Results

### Model of mutation-selection-migration and genetic drift

To quantify the conditions under which between-host transmission is expected to affect the spectrum of mutations shared across hosts, we used a model of mutation-selection-migration and genetic drift (see Materials and Methods). The simulations predict that when the number of migrants is >>1, the metapopulation behaves as a single population and all beneficial mutations are shared across demes (**fig. 1A and supplementary fig. S1**). At the other extreme, when the number of migrants is extremely small, demes behave as close to independent populations and the level of shared adaptive polymorphisms is expected to be very low.

**Fig. 1.**
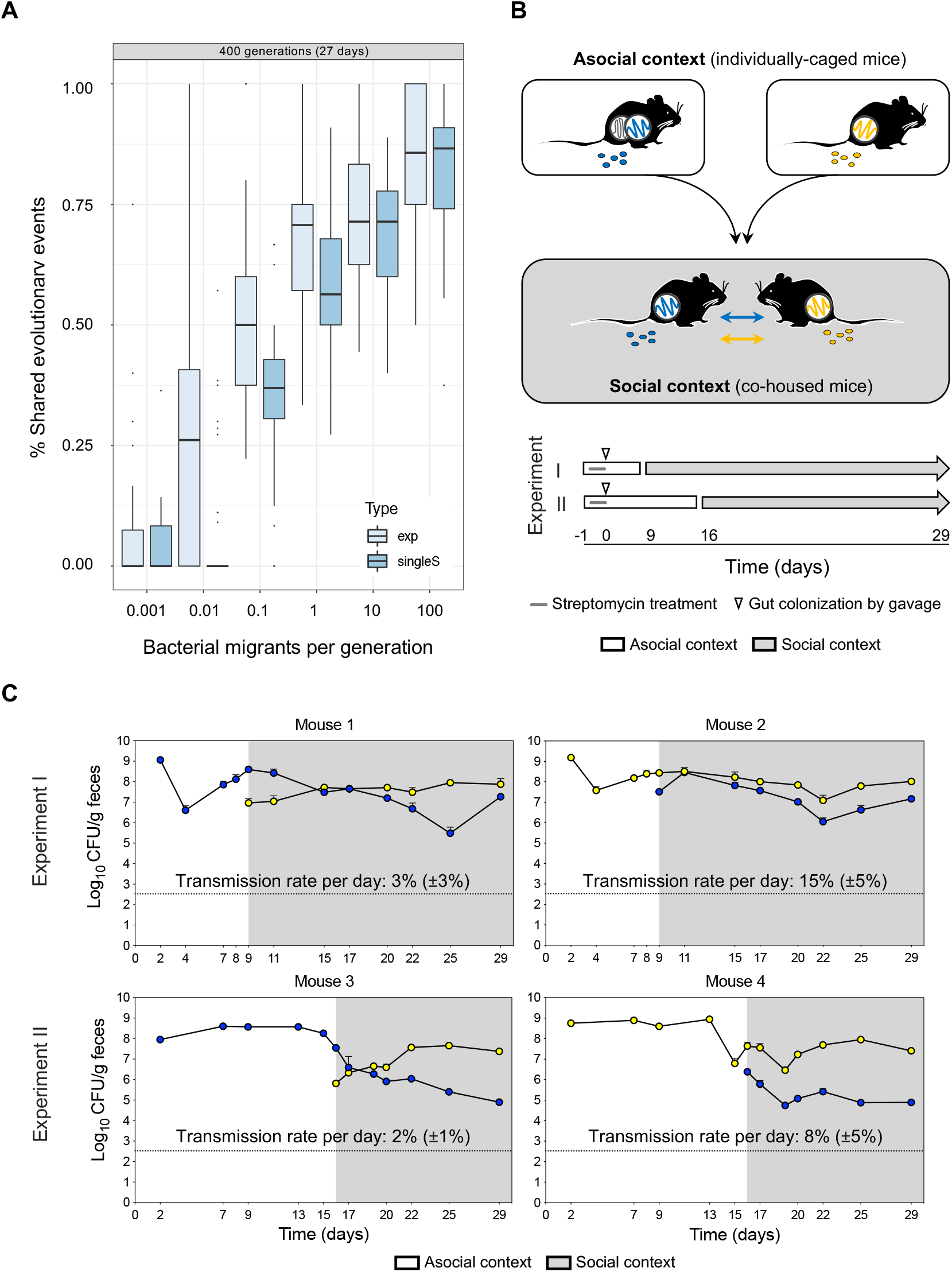
*E. coli* transmission between mice. (**A**) Model of bacterial transmission. The relation between the level of shared evolutionary events and migration in a model where all mutations have the same effect (type singleS) and in one where mutations are exponentially distributed (type exp). Parameter values *N*=10000, *U*_*d*_=0.01; *U*_*a*_=0.00001; *S*_*d*_=0.1; *S*_*a*_=0.1; Populations accumulate mutations during 400 generations, in a time scale of *E. coli* evolution in the gut of 27 days. (**B**) Experimental design to estimate migration rates. Mice were colonized with an *E. coli* clone expressing either a cyan (CFP) or a yellow (YFP) fluorescent protein (10^7^ CFUs) while in an asocial environment (individually caging). In experiment I, mice (*n* = 2) were moved to a social environment (co-housing) at day 8, while in experiment II (*n* = 2), mice were moved to a social environment at day 15. (**C**) Fecal CFP and YFP *E. coli* loads were assessed under asocial and social contexts. Gray color indicates mice in a social context (co-housing). Error bars represent 2X standard error (2SE) and the dashed line indicates the limit of detection (330 CFU/g of feces). *E. coli* transmission rate (%): percentage of *E. coli* cells in the mouse gut that resulted from transmission events, each day, from the remaining mouse inside the cage. Error represent 2X standard error (2SE). See Table 2.

### High levels of strain transmission under co-housing

In the simplest theoretical model homogenization is expected if bacterial transmission is high, with pervasive sharing of mutations between hosts. To be able to test this hypothesis that transmission increases shared polymorphism and accelerates adaptation, we first quantified the extent of migration across hosts using two isogenic *E. coli* strains (**Table 1**), each carrying a different neutral marker: yellow fluorescent protein (YFP) or cyan fluorescent protein (CFP). The experimental design is described in **fig. 1B**: each mouse was colonized with one of the labeled lineages, CFP or YFP during a period of *T* days (*T* = 7 or 14 days, **fig. 1B** - Experiments I and II, respectively). After this period, the mice were co-housed (social regime) to allow for microbiome transmission between hosts and to quantify the rate of transmission of *E. coli* between hosts. The co-housing after two weeks of colonization was performed to test if transmission would still occur after a period when within-host adaptation could already have occurred. We found transmission of the *E. coli* across mice to be pervasive and extremely fast (**fig. 1C, Table 2**), with co-colonization with both YFP and CFP lineages occurring in less than 24 h of co-housing both animals. On average, 1.5 × 10^7^ *E. coli* cells [±1 × 10^7^, 2x standard error (2SE)] transmit to a new mouse per day, assuming that migration happens at a constant rate (**Table 2**). We thus estimate a rate of transmission of 7% (± 3% 2SE) per day for this *E. coli* strain (**fig. 2C, Table 2**). This rate is similar to that observed in germ-free mice (Vasquez et al. 2021). This establishes that co-housing constitutes a regime of high migration and thus theoretically expected to result in high levels of allele sharing (**fig. 1A**).

**Fig. 2.**
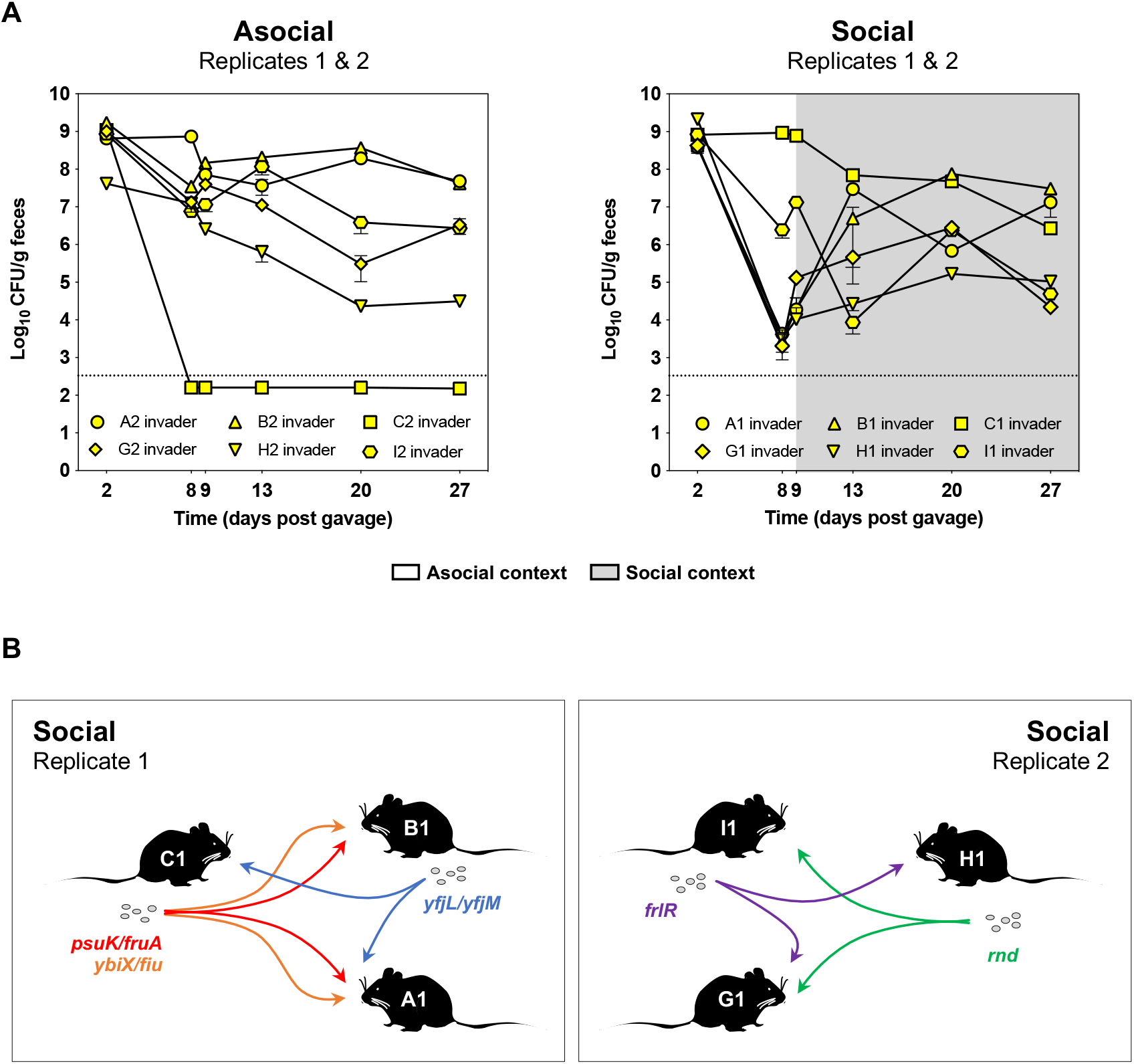
*Escherichia coli* evolution while transmitting among multiple hosts (social context). (**A**) Loads of invader *E. coli* clone introduced by gavage into the mice in an asocial or social regimes. Error bars represent (2SE). The limit of detection (dashed line) is 330 CFU/g of feces. (**B**) Transmission of *E. coli* with specific mutations (*e*.*g*., *psuK/fruA, ybi/fiu, yjfL/yjfM, frlR* and *rnd*) from donor to recipient mice. See Table 6.

### Rate of molecular evolution under co-housing

Having established that bacterial migration is common in co-housed mice, we designed a new experimental setting (**supplementary fig. S2**) to test for the influence of transmission in the patterns of molecular evolution. In 5 out of 6 mice living asocially successful colonization with the invader *E. coli* was observed, whereas under social conditions all mice were successfully colonized (**fig. 2A, Table 3)**. Provided that colonization was successful for at least one month (27 days), we did not find significant differences in the loads of the invader *E. coli* in the asocial *versus* social environments, stabilizing around log10 6.3 (±0.58, 2SE) CFU/g feces (**fig. 2A and supplementary fig. S3**).

We then sequenced pools of clones that evolved in the mouse gut for either a month (27 days) or more than 3 months (104 days) using Illumina technology and mapped the reads to the genome of their ancestor (**Table 4)**. Parallel mutations, a sign of adaptive evolution, were noticed as previously (Barroso-Batista et al. 2014; Frazão et al. 2019; Barroso-Batista, Pedro, Sales-Dias, Catarina J.G. Pinto, et al. 2020; Frazão et al. 2022), being present in both the *E. coli* populations isolated from animals in social and asocial settings (**Table 5**). Adaptive mutations, such as *psuK/fruA* and *frlR*, could be traced back by sequencing populations from previous sampling points. This allowed us to identify the mice where the adaptive mutations were selected originally and then transmitted to the remaining mice in the same cage (**fig. 2B and Table 6**). No synonymous mutations were detected in clones sampled from the mice in a social setting, leading to a ratio of non-synonymous to synonymous mutations dN/dS infinity 1 in all the animals. In populations isolated from individually caged mice the dN/dS ratio was > 1 in only 3/5 animals **(Table 5)**, suggesting that adaptive evolution is stronger when bacterial transmission is occurring.

We then quantified how the dynamics of mutations accumulation in each host is affected by its social environment. We calculated the sum of the allele frequencies at each sampling point along time (M(t)) as a measure of the rate of molecular evolution that *E. coli* undergoes when colonizing in a social *versus* asocial setting **(fig. 3A and Table 5)**. Mutations accumulated at an average rate of 2.8×10^−3^ (±5.9×10^−4^ 2SE) or 2.9×10^−3^ (±9.3×10^−4^ 2SE) per genome per generation, in a social or asocial regime, respectively **(fig. 3B and Table 5)**, suggesting that the social regime did not affect the rate of molecular evolution.

**Fig. 3.**
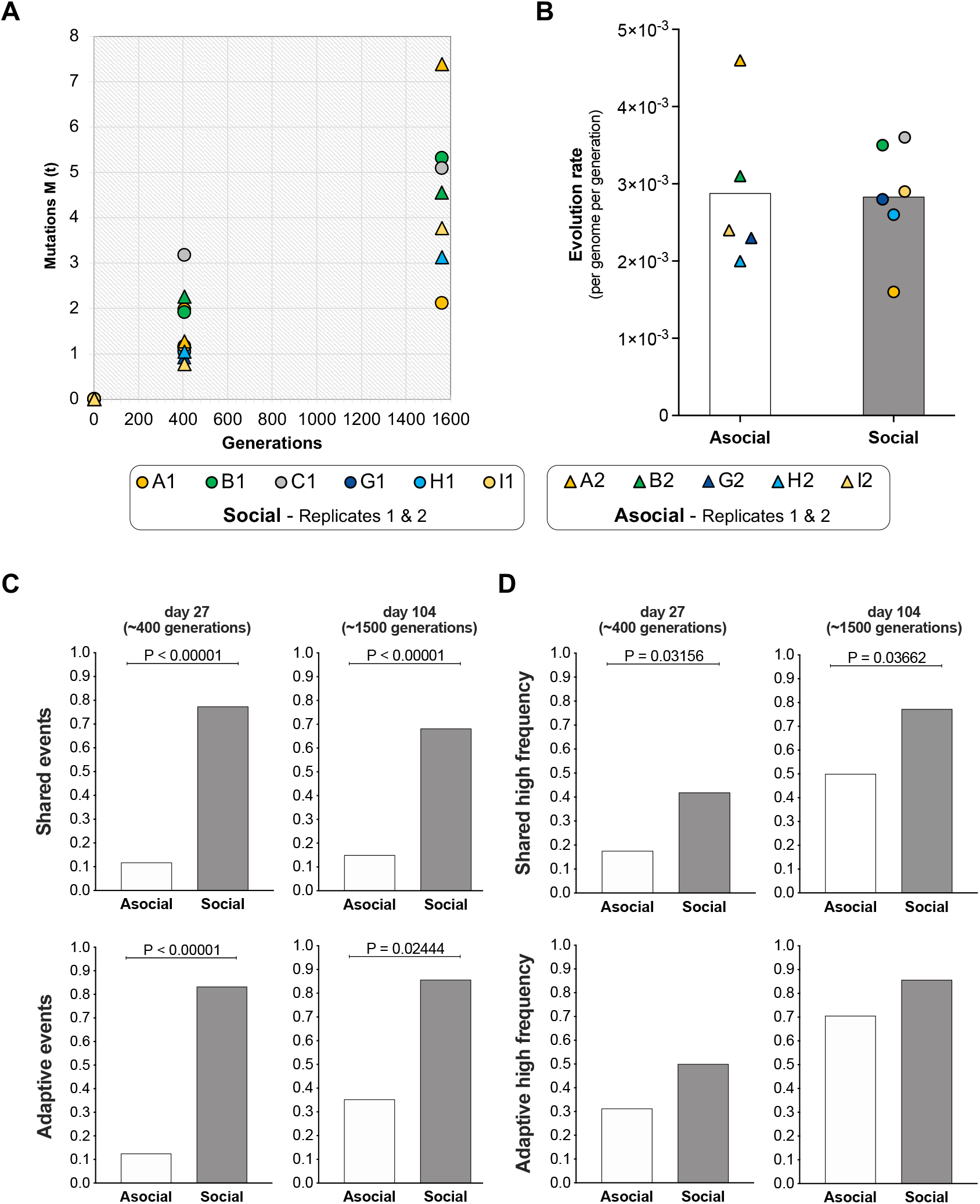
Molecular *E. coli* evolution in a social and asocial context. (**A**) *E. coli* mutation accumulation during *in vivo* evolution. Mutations M(t) correspond to the sum of allele frequencies at each sampling point along time (generations). (**B**) *E. coli* evolution rate based on the M(t). The slope (intersect = 0) of the mutations M(t) corresponds to the *E. coli* evolution rate, per genome, per generation in each mouse from each biological replicate (social and asocial regimes). See Table 5. Biological replicate 1 includes mice: A1, B1 and C1 (social regime) and A2 and B2 (asocial regime). Biological replicate 2 includes mice: G1, H1 and I1 (social regime) and G2, H2 and I2 (asocial regime). Bars represent the average rate of evolution in social or asocial regimes. (**C**) Proportion of shared or adaptive evolutionary events observed in the invader *E. coli* populations isolated from mice living in an asocial or social regime at day 27 or 104 (**D**). Proportion of shared or adaptive high frequency evolutionary events (>50%) observed in the invader *E. coli* populations at day 27 or 104. Statistics correspond to Binomial Test for proportions.

### A social regime boosts sharing of evolutionary events

The number of mutations accumulated per mouse was marginally lower in the social environment (median 3.5 in the social *versus* 6 in the asocial, Mann-Whitney two tailed test U=5, p=0.08, **Table 4**). In the first cohort of co-housed mice, 9 out of 12 (75%) mutations were common across all 3 mice, whereas in the second cohort 6 out of 8 (75%) were common. In the asocial context, only 4 out of 16 (25%) mutations were common across all mice in the first cohort, while none was common in the second (**Table 4**). Consistent with the presence of a resident *E. coli* colonizing the mouse gut (Frazão et al. 2019) (**supplementary fig. S4**), 15 phage-driven HGT events were detected during the same time period (27 days: ~400 generations). Of these, 11 occurred in co-housed animals (social regime, n=6 mice), while only 4 events were observed in individually-caged animals (asocial regime, n=5 mice), as the resident strain was present only in 2 out of the 5 animals (**supplementary fig. S4**). In the first and second cohorts of cohoused mice, 100% of phage-driven HGT events were common across all animals, while in the asocial context phage-driven HGT events were observed only in 2 animals where the resident was co-colonizing (**Table 7**).

We next evaluated how many evolutionary events (mutations and/or phage-driven HGT) are shared between individuals. The proportion of evolutionary events (mutation and phage-driven HGT) shared among hosts at day 27 (~400 generations) post-colonization is significantly higher under a social environment — 77.4% (24 shared out of 31 evolutionary events) — than in an asocial regime —11.8% (4 out of 34 events) (Binomial Test for proportions, *P*<0.00001; **fig. 3C and Table 4, 7 and 8)**. We next asked whether the level of allele sharing in co-housed mice would still be higher after long-term evolution and sequenced pools of *E. coli* clones evolved for 104 days (~1500 generations) in the first cohort of mice. The sharing of evolutionary events was still significantly higher under co-housing (68.2%) than in the asocial environment, (15%), even after more than a thousand generations of evolution (Binomial Test for proportions, *P*<0.00001; **fig. 3C and Table 4, 7 and 8**). Importantly, when focusing on *bona fide* adaptive evolutionary events, *i*.*e*., those that were observed in more than one independent mouse, among the two social regimes, the same trend towards more genetic parallelism in the social regime was observed (**fig. 3C**). The proportion of adaptive events (mutations and phage-driven HGT) that were shared among the hosts at day 27 (~400 generations) was significantly higher under a social environment, 83.2% (15 out of 18 evolutionary events) than in an asocial regime, 12.5% (2 out of 16 events) (Binomial Test for proportions, *P*<0.00001; **fig. 3C, Table 7, 9 and 10)**. Additionally, even after 104 days (~1500 generations), the proportion of adaptive events was significantly higher, 85.7%, in animals under a social regime than in asocial conditions, 35.3% (Binomial Test for proportions, *P*=0.02444; **fig. 3C, Table 7, 9 and 10)**.

### A social regime increases the proportion of high frequency evolutionary events

To understand if the social regime affects the frequency trajectory of the evolutionary events, we started by analyzing the proportion of high frequency events (> 50%) among the total events observed in *E. coli* populations isolated from mice living in the social or asocial regimes. The proportion of total mutations or HGT events that reached high frequency (**Fig. 3D**, frequencies > 50%) was significantly higher at day 27 (~400 generations) when *E. coli* colonized mice living in a social regime, 41.9%, than in animals independently caged, 17.6% (Binomial Test for proportions, *P*=0.03156; **fig. 3D, Table 8**). At day 104 (~1500 generations) the proportion of high frequency events (**fig. 3D**) remained high in the co-housed animals, 77.3%, while in mice living in independent cages it was only 50% (Binomial Test for proportions, *P*=0.03662; **fig. 3D, Table 8**). When the same analysis was performed focusing only on *bona fide* adaptive events (mutations and/or phage-driven HGT events observed in more than one independent animal) the proportion of high frequency events (**fig. 3D, Table 10**) was not significantly different between *E. coli* populations isolated from mice in social or asocial conditions. This could be due to lack of statistical power as we observed a limited number of mutational events when compared with the total of events. Nevertheless, adaptive events show the same tendency as when considering all evolutionary events, *i*.*e*., a higher number of high frequency events in the social regime (**fig. 3D, Table 10**).

## Discussion

Bacterial evolution in the mammalian intestine has been studied mainly under asocial conditions, where bacterial transmission among mammalian healthy hosts does not take place (Barroso-Batista et al. 2014; Barroso-Batista et al. 2015; Lourenço et al. 2016; Lescat et al. 2017; Sousa et al. 2017; Frazão et al. 2019; Ghalayini et al. 2019; Ramiro et al. 2020; Frazão et al. 2022). To our knowledge, the evolution of *E. coli* in the presence *versus* absence of bacterial transmission in mice bearing a microbiota is compared here for the first time. We established a mouse model of bacterial transmission to compare *E. coli* evolution when colonizing the gut of mice living in a social or asocial regime. We found that in mice inhabiting the same cage 7% (±3% 2SE) of the *E. coli* colonizing each mouse gut results from daily migration events. It is known that *E. coli* can adapt to better colonize the mouse intestine in less than a week of colonization, with the mutation and phage-driven HGT events occurring concomitantly (Frazão et al. 2019). Here we observed that *E. coli* transmission between hosts can still occur even when an adapted *E. coli* is already colonizing the gut. Interestingly, the first clone to colonize a given mouse was not always the one ending up as dominant after a month of evolution, suggesting that priority effects (Sprockett et al. 2018) should not be dominant in this species.

The *E. coli* population size, a key aspect in determining the level of variability in a population and the effectiveness of selection relative to drift (Charlesworth 2009), was not affected by transmission, with populations isolated from mice living socially or asocially presenting similar loads. The *E. coli* cells transmitting among microbiome-bearing mice was comparable to the 10% migration inferred in a study using germ-free animals (Vasquez et al. 2021), suggesting that *E. coli* transmits between hosts at a very similar rate independently of the presence or absence of a microbiome. Interestingly, we show invasion and coexistence of the same strain of *E. coli* (expressing different fluorescent markers) in mice living in a social regime, suggesting that strain colonization resistance, observed for other important species in the gut such as *Bacteroides fragilis* (Lee et al. 2013), is absent from the *E. coli* species.

Importantly, whole-genome sequencing of populations revealed that when in a social regime, where host-to-host bacterial transmission is pervasive, the evolution of *E. coli* is always characterized by an elevated ratio of non-synonymous to synonymous mutations, indicative of evolution driven by strong selection of adaptive haplotypes, a phenomenon we found to be less common in an asocial regime.

Our theoretical model of bacterial transmission predicted that the outcome of bacterial migration between hosts would affect the evolutionary process in terms of the number of shared evolutionary events between hosts. The model simulations predicted that when the number of migrant clones between mice is large enough (>>1), the metapopulation of co-housed mice acts as a single population and genetic changes are shared across hosts. Alternatively, when transmitting a very low number of bacterial migrants, each host behaves as an independent population, with the level of sharing of genetic evolutionary events being extremely low. Indeed, a significantly higher proportion of shared evolutionary events (total and adaptive) was found in *E. coli* populations from the social condition. These observations relate not only to the mutational process, but also to the evolutionary phage-driven events. We were able to study the latter process because our mouse cohort was colonized with a resident *E. coli* carrying several active prophages that can be transferred to the invader strain, thus conferring an adaptive potential to consume sugars that are common in the gut (Frazão et al. 2019). The adaptive mutational process targeted mostly the same biological processes in both invader *E. coli* populations, isolated from the social or asocial regime, namely production of pyrimidine nucleotides from pseudouridine (*psuK/fruA* mutation) and fructoselysin consumption (*frlR* mutation).

Phages are known key players in shaping the composition and diversity of bacterial communities in many environments, namely in the gut of mice and humans (Kim and Bae 2018; Frazão et al. 2019; Sutton and Hill 2019). Here we observed that phage-driven HGT events, namely the phages Nef and KingRac were transferred to the invader *E. coli* genome as observed previously (Frazão et al. 2019). This process was independent of the social or asocial condition, though limited to when co-colonization with both the resident and invader lineages occurred. Nevertheless, under a social regime, bacterial transmission across hosts spread these adaptive phage-driven events across all co-housed animals.

We found that under the social regime total evolutionary events reached significantly higher frequencies in the invader population, with adaptive events following the same tendency. Invader populations evolving during transmission did not show any signs of diversifying selection. On the contrary, these populations exhibited genetic homogeneity and an elevated dN/dS ratio indicative of adaptive evolution driven by strong selection, which was indeed predicted in the simulations modeling selection under strong migration.

The present study also reveals that the evolution rate of *E. coli* is unchanged in both social and asocial settings, being on average 2.9×10^−3^ (±5×10^−4^ 2SE) mutations/genome/generation. Interestingly, this rate is similar to those reported in previous studies *in vitro* (Good et al. 2017) and *in vivo* (Frazão et al. 2022), where bacterial host-to-host transmission was not taking place. Thus, as proposed previously (Frazão et al. 2022), *E. coli* appears to follow a clock-like rate of evolution, which the present data show is independent of the host social context where *E. coli* evolves.

We conclude that the effect of migration should be taken into account when analyzing the evolutionary history of a bacterium colonizing the mammalian gut. Our findings demonstrate that the evolution of new bacterial strains invading the gut microbiota can be strongly affected by the host’s social environment, which calls for further experimental evolution studies comparing social *versus* asocial conditions.

## Materials and Methods

### *Escherichia coli* clones

The clone used in the present study is described in Table 1. Ancestral and evolved *E. coli* clones were grown at 37°C under aeration in brain heart infusion (BHI) or Luria broth (LB). Media were supplemented with antibiotics when specified. Serial plating of 1X PBS dilutions of feces in LB agar plates supplemented with the appropriate antibiotics were incubated overnight and YFP-labeled bacterial numbers were assessed by counting the fluorescent colonies using a fluorescent stereoscope (SteREO Lumar, Carl Zeiss) with the detection limit for bacterial plating being 330 CFU/g of feces.

### *In vivo E. coli* evolution experiment

Mice drank water with streptomycin (5 g/L) for 24 h before a 4-h starvation period of food and water. The animals were then inoculated by gavage with 100 µL of an invader *E. coli* bacterial suspension of ~10^8^ colony forming units (CFUs). Mice littermates A1/A2, B1/B2, C1/C2, G1/G2, H1/H2 and I1/I2 were all gavaged and successfully gut colonized with the invader *E. coli* clone, except mouse C2. The same letter describes animals that are littermates, while number 1 or 2 means the mice are in a social or asocial regime, respectively. Five (A2, B2, G2, H2 and I2) out of the 11 mice colonized with *E. coli* were already followed, in previous studies (Frazão et al. 2019; Frazão et al. 2022). Six- to eight-week-old C57BL/6J female mice were kept in ventilated cages under specified pathogen free (SPF) barrier conditions at the Instituto Gulbenkian de Ciência (IGC) animal facility. The gut microbiota of mice used in the experiments is natural to the animals and not the result of the introduction of a defined microbiota into germ-free animals. Fecal pellets were collected and stored in 15% glycerol at −80°C for later analysis. This research project was ethically reviewed and approved by the Ethics Committee of the Instituto Gulbenkian de Ciência (license reference: A009.2010) and by the Portuguese National Entity that regulates the use of laboratory animals (DGAV – Direção Geral de Alimentação e Veterinária; license reference: 008958). All experiments conducted on animals followed the Portuguese (Decreto-Lei nº 113/2013) and European (Directive 2010/63/EU) legislations concerning housing, husbandry and animal welfare.

### Whole-genome sequencing of evolved *E. coli* populations

DNA was extracted (40) from *E. coli* populations (mixture of >1000 clones) growing in LB plates supplemented with antibiotic to avoid contamination. DNA concentration and purity were quantified using Qubit and NanoDrop, respectively. DNA library construction and sequencing were carried out by the IGC Genomics facility using the Illumina platform. The mean coverage per population was 200x and processing of raw reads and variants analysis was performed as previously described (Frazão et al. 2019; Frazão et al. 2022).

### Model of bacterial transmission

A model of mutation-selection-migration and genetic drift was used to predict the effect of bacterial transmission on the landscape of mutations shared among populations of *E. coli* colonizing the gut of mice. For simplicity we assume that there are two demes of equal carrying capacity (*N*). Individuals reproduce clonally within each deme and acquire deleterious mutations at a rate *Ud* and fitness increasing mutations (adaptive mutations) at a rate *U*_*a*_ per generation (typically 1/100 of *U*_*d*_, (Perfeito et al. 2007)). Under the simplest model, the effect of a deleterious mutation is assumed to be constant (*S*_*d*_) as well as that of an advantageous mutation (*S*_*a*_). A distribution of fitness effects, where the effects of both mutations are taken from exponential distributions of mean *S*_*d*_ or *S*_*a*_, is also modeled to establish the generality of the qualitative expectations. Selection acts within each deme and *N* individuals are chosen according to their fitness. The model ignores any trade-offs of mutation effects outside the host environment, implicitly assuming transmission by direct contact. Migration follows selection, where a Poisson number of migrants (*Nm*) is exchanged between two demes according to the migration rate *m*. To measure the level of shared adaptations across demes, each individual adaptive portion of the genome is explicitly modeled with a finite size *G* and a bi-allelic model for adaptive mutations. Thus, the mutation rate towards new advantageous alleles is *Ua*/*G* per site per generation. Deleterious mutations follow an infinite site model, as these are in principle much more numerous than beneficial mutations.

### Statistical analysis

A linear mixed-effects model was used to do the analysis of temporal load differences of the invader *E. coli* from days 9 to 27, with time as repeated measures and housing as treatment factor. A Mann-Whitney test was used to compare the evolution rate in social and asocial groups. A Binomial Test was used to compare proportions in the two regimes (social and asocial). A *P*-value < 0.05 was considered for statistical significance.

## Data and Resource Availability

Raw sequencing reads were deposited in the sequence read archive under the name of bioproject PRJNA930727.

## Acknowledgements

We thank the Evolutionary Biology group for helpful discussions. We would also like to thank the personnel of the IGC’s Rodent Facility, Genomic Facility and the Bioinformatics Unit for their assistance. N.F. was supported by “Fundação para a Ciência e Tecnologia” (FCT), fellowship SFRH/BPD/11075/2015. This work was also supported by project Global Gut Health Nature Research/Yakult Grant 623877 and ONEIDA project (LISBOA-01-0145-FEDER-016417) co-funded by FEEI – “Fundos Europeus Estruturais e de Investimento” from “Programa Operacional Regional Lisboa 2020”, by national funds from FCT and FCT Project PTDC/BIA-EVL/7546/2020.

## Contributions

I.G. and N.F. designed and coordinated the study.

N.F. performed the experiments. N.F. and I.G. analyzed the results and wrote the manuscript giving final approval for publication.

## Supplementary Figures

**Figure S1.**
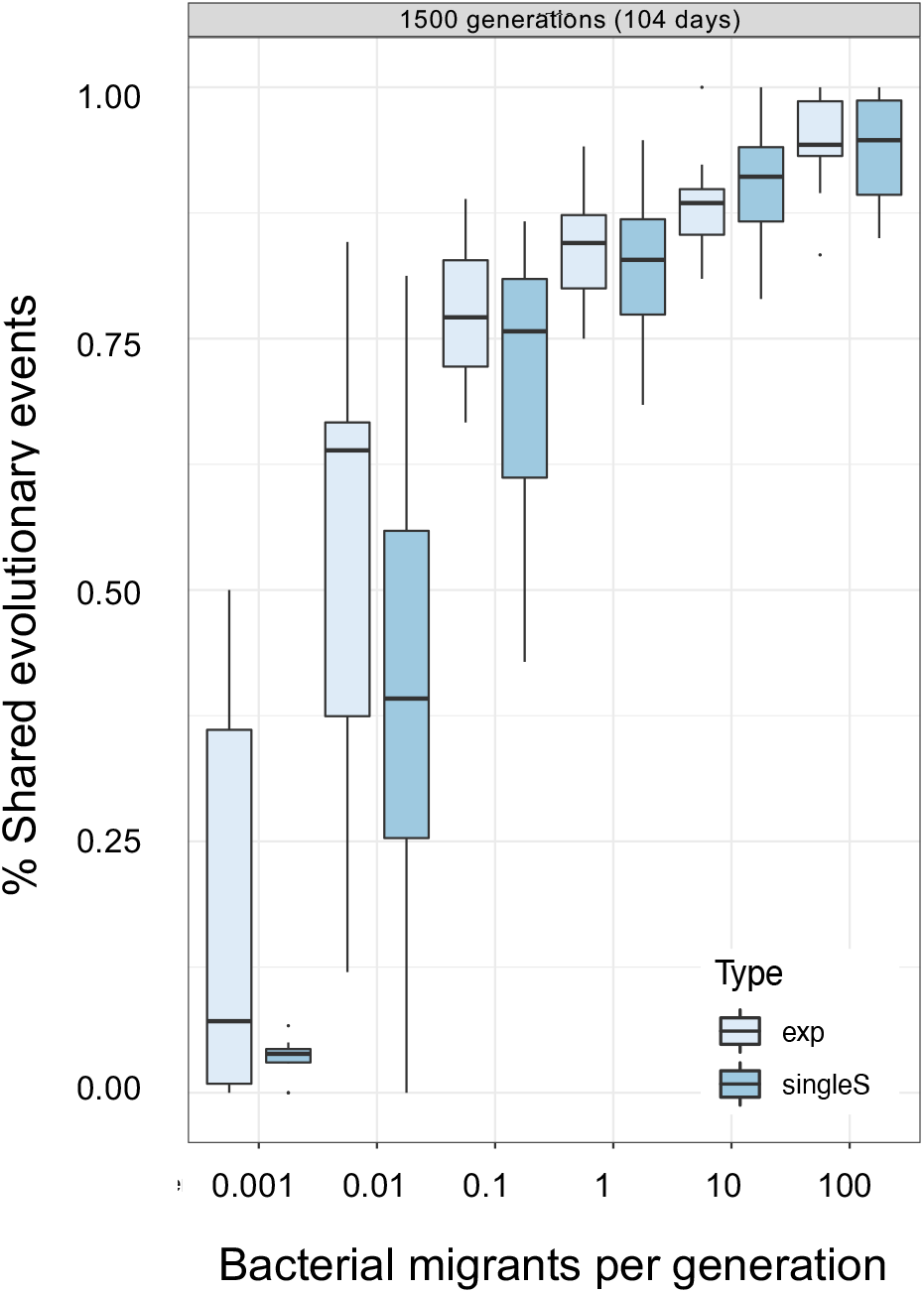
Model of bacterial transmission. The relation between the level of shared evolutionary events and migration in a model where all mutations have the same effect (type singleS) and in one where mutations are exponentially distributed (type exp). Parameter values *N*=10000, *U*_*d*_=0.01; *U*_*a*_=0.00001; *S*_*d*_=0.1; *S*_*a*_=0.1; Populations accumulate mutations during 1500 generations, in a time scale of *E. coli* evolution in the gut of 104 days.

**Figure S2.**
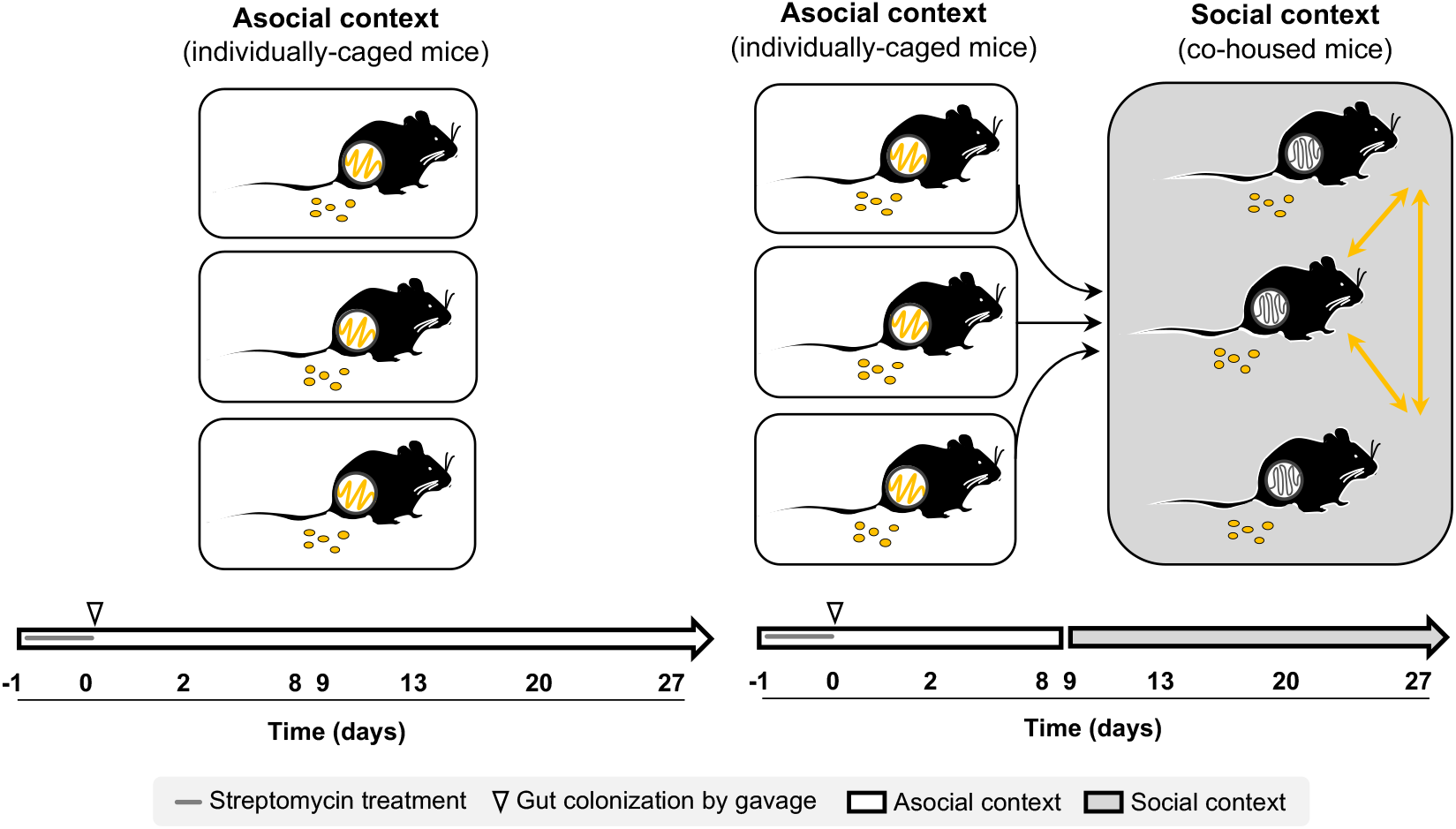
Experimental setup to study *Escherichia coli* evolution while transmitting among multiple hosts (social context). Adult female C57BL/6J mice (6-8 weeks old) from different litters were treated with streptomycin (5 g/L) for 24 h and then gavaged with 10^7^ CFU/mL of the *E. coli gatZ* clone 5 expressing yellow fluorescent protein (YFP). After gavage, fecal pellets were sampled in order to analyze the *E. coli* load in mice feces that were under either an asocial or social context. Gray color background means that the feces sampling was performed with mice in a social context (co-housed).

**Figure S3.**
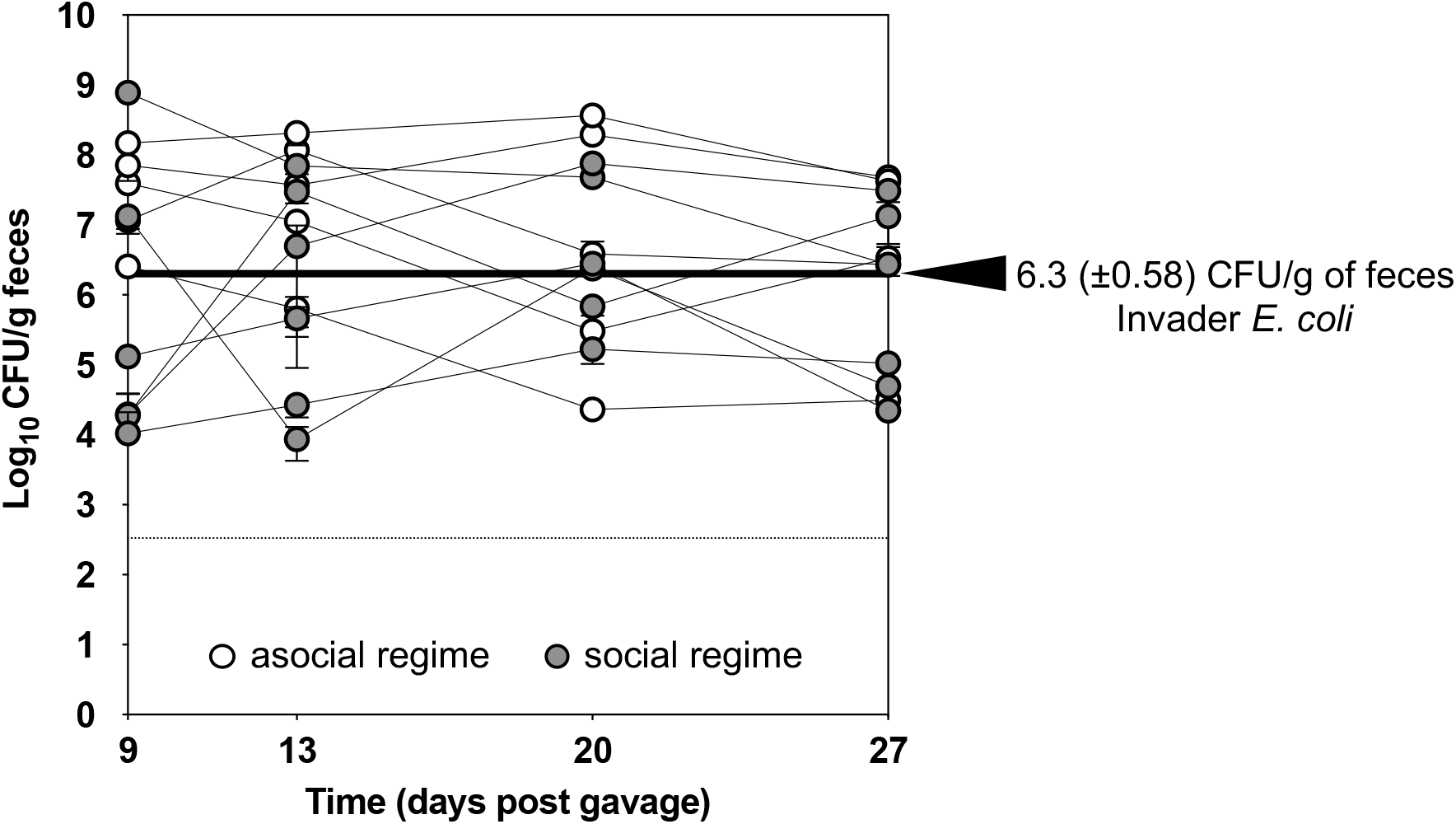
Invader *E. coli* loads in social and associal settings. Using a linear mixed effects model analysis of the temporal loads we were able to quantify the average invader *E. coli* load (black arrow) when colonizing the gut of mice. The linear mixed effects model had into account the invader *E. coli* loads in mice A2, B2, G2, H2 and I2 (white circles, asocial regime), and mice A1, B1, C1, G1, H1 and I1 (gray circles, social regime). No significant difference across time was found either between days or between mice living in social or asocial regimes regarding the invader *E. coli* loads. The dotted line indicates the detection limit (330 CFU/g of feces).

**Figure S4.**
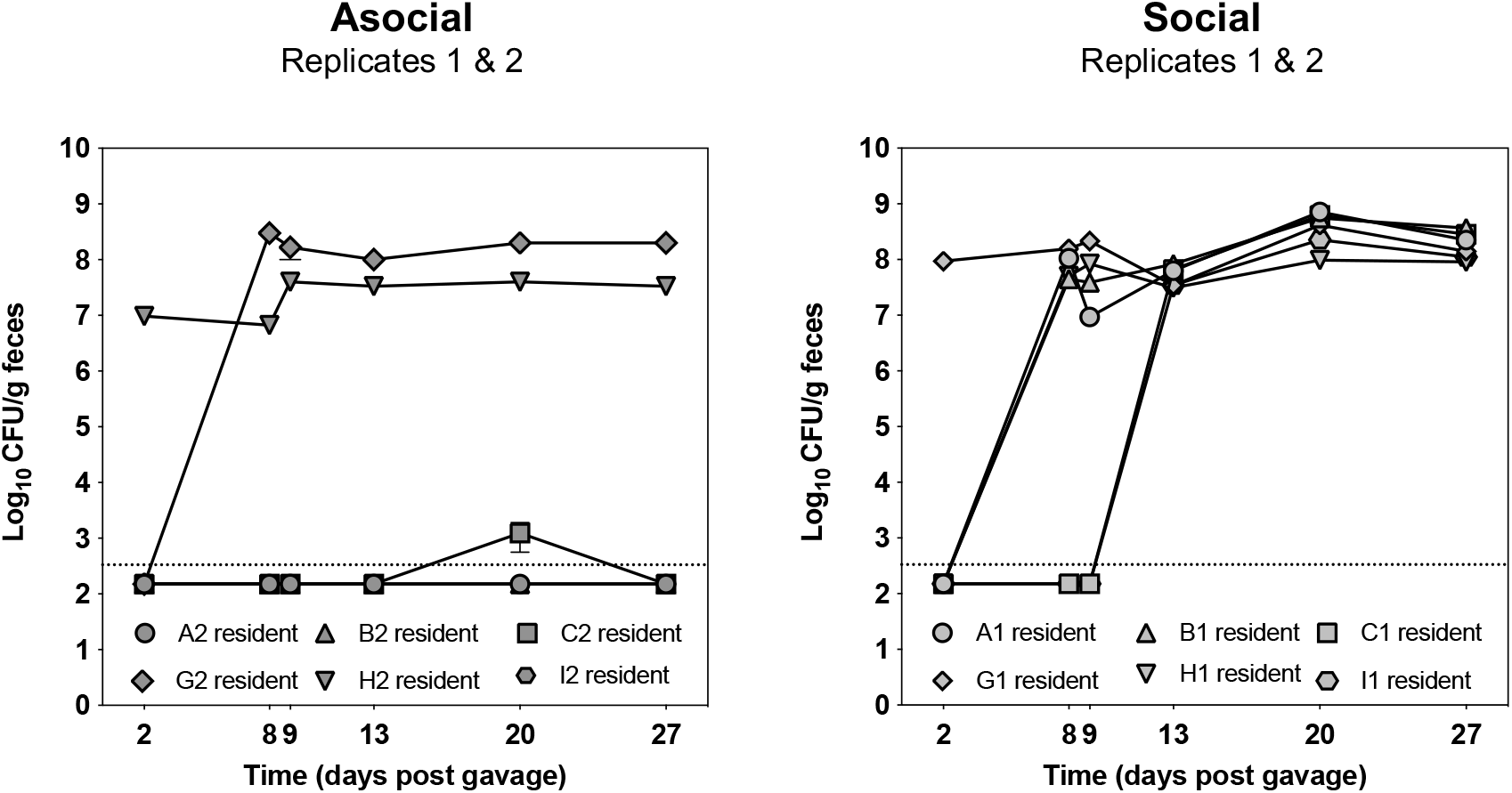
Resident *Escherichia coli* colonizing mice in social *versus* asocial regimes. Loads of the Resident *E. coli* colonizing the mice in an asocial or social regimes. Error bars represent (2SE). The limit of detection (dashed line) is 330 CFU/g of feces.

